# Identification of genes with enriched expression in early developing mouse cone photoreceptors

**DOI:** 10.1101/538611

**Authors:** Diego F. Buenaventura, Adrianne Corseri, Mark M. Emerson

**Affiliations:** Department of Biology, The City College of New York, City University of New York, New York, NY, 10031; Biology Ph.D. Program, Graduate Center, City University of New York, New York, NY, 10031

**Author notes:** Department of Biology, New York University, New York, NY 10003.

## Abstract

Cone photoreceptors are the critical first cells that mediate high acuity vision. Despite their importance and their potential use in cell-based therapies for retinal diseases, there is a lack of knowledge about the early developmental stages of these cells. Here we characterize the expression of the homeobox transcription factor Lhx4 as an early and enriched cone photoreceptor expressed gene in both chicken and mouse. A Lhx4 GFP reporter mouse was found to recapitulate this early cone photoreceptor expression and was used to purify and profile embryonic mouse cone photoreceptors by single cell RNA sequencing. This enrichment in cone photoreceptors allowed for the robust identification of genes associated with the early cone transcriptome and also identified subpopulations of these cells. A comparison to previously reported datasets allowed the classification of genes according to developmental timing, cell type specificity, and whether they were regulated by the rod transcription factor Nrl. This analysis has extended the set of known early cone enriched genes and identified those that are regulated independently of Nrl. This report furthers our knowledge of the transcriptional events that occur in early cone photoreceptors.

## INTRODUCTION

Cone and rod photoreceptors are the photosensitive cells of the retina that contribute to image formation. Cones mediate color discrimination and high acuity vision while rods provide photosensitivity in low-light conditions. Given the importance of cones in high acuity and color vision, deficiency in this cell type as a result of conditions such as retinitis pigmentosa or macular degeneration, lead to a debilitating loss of vision^1^. As such, the development of cell-based therapeutic strategies based on the formation of new cone photoreceptors is a promising strategy. ^2^ However, there is presently a gap in our knowledge of the gene regulatory networks that control the genesis of these cells as well as the early steps in their differentiation. Thus, the design of informed strategies to direct cone production and how to appropriately benchmark the differentiation of de novo generated cells is lacking.

Two main strategies have been used to investigate the early gene regulation programs of cone photoreceptors. One has been to develop reagents to label developing cone cells or the RPCs that generate them and use high-throughput methods to identify the genes with enriched expression in these cells compared to other cells present at the time ^3, 4^. Such methods have also been used at later differentiation timepoints ^5–7^. While these methods provide critical information, they also rely on transcriptional reporters that may have expression that is broader than just cone photoreceptors and do not provide cellular resolution of these gene expression patterns. The second strategy to examine cone photoreceptor gene expression has been through the use of the Nrl mouse model. Nrl is a critical rod-expressed transcription factor that is necessary to promote rod gene expression and repress cone gene expression in rod cells ^8^. The Nrl mouse knockout leads to a large increase in cone gene expression as the large number of rods in the mouse undergo derepression of cone genes^9–11^. These Nrl knockout rods have been interpreted as either a complete fate switch to cones or a partial conversion to “cods”^8, 12, 13^. As these cells have been used in a number of studies to model cone photoreceptors, the extent to which these cells are transformed to the cone fate is important for both a justification in using them as a model for endogenous cone cells and to understand photoreceptor diversification.

Here, we identified the Lim Homeobox Protein 4 (LHX4) gene as enriched in cones during early chick retinal development, in addition to bipolar cells. In the mouse, LHX4 was also determined to be a reliable marker for cones during early retinal development but with expanded expression in late embryonic development and eventual expression in BCs. A LHX4::GFP transgenic line ^14^ recapitulated the endogenous LHX4 expression pattern and was used to generate a single cell dataset highly enriched in cone photoreceptors, which provided an in-depth look at the molecular profile of these cells in the earliest stages of differentiation in the mouse. A comparison was made between previous datasets that targeted mammalian cones and photoreceptors, including those made in the Nrl knockout mouse. This led to the identification of unique cone expression signatures not observed in previous datasets, including those of Nrl knockout rods, supporting previous observations that these cells are not completely transformed into cones.

## RESULTS

### LHX4 is present in early cone photoreceptors in the chicken retina

Recently, we established the transcriptional profile of retinal progenitor cells (RPCs), defined by the activity of the ThrbCRM1 element, that are biased towards the cone and horizontal cell (HC) fate in the early chick retina^3^. Using this dataset, we screened for potential cone-enriched transcripts that could serve as markers for early cones. After establishing a criterion for >1.5-fold change score between cone/HC RPCs and other concurrent populations (enriched in “Other early retinal progenitors”) we selected for transcription factors (TFs) enriched in the cone/HC RPCs (Fig 1 A). We identified the LIM homeobox 4 (LHX4) gene as highly enriched, along with known TFs in this population such as THRB, ONECUT1, and OTX2. This transcript has significant fold change (b = 3.3) and a low number of reads in the non-ThrbCRM1 active population, suggesting high specificity towards the cone/HC RPC population at this time (Fig 1 B).

**Figure 1.**
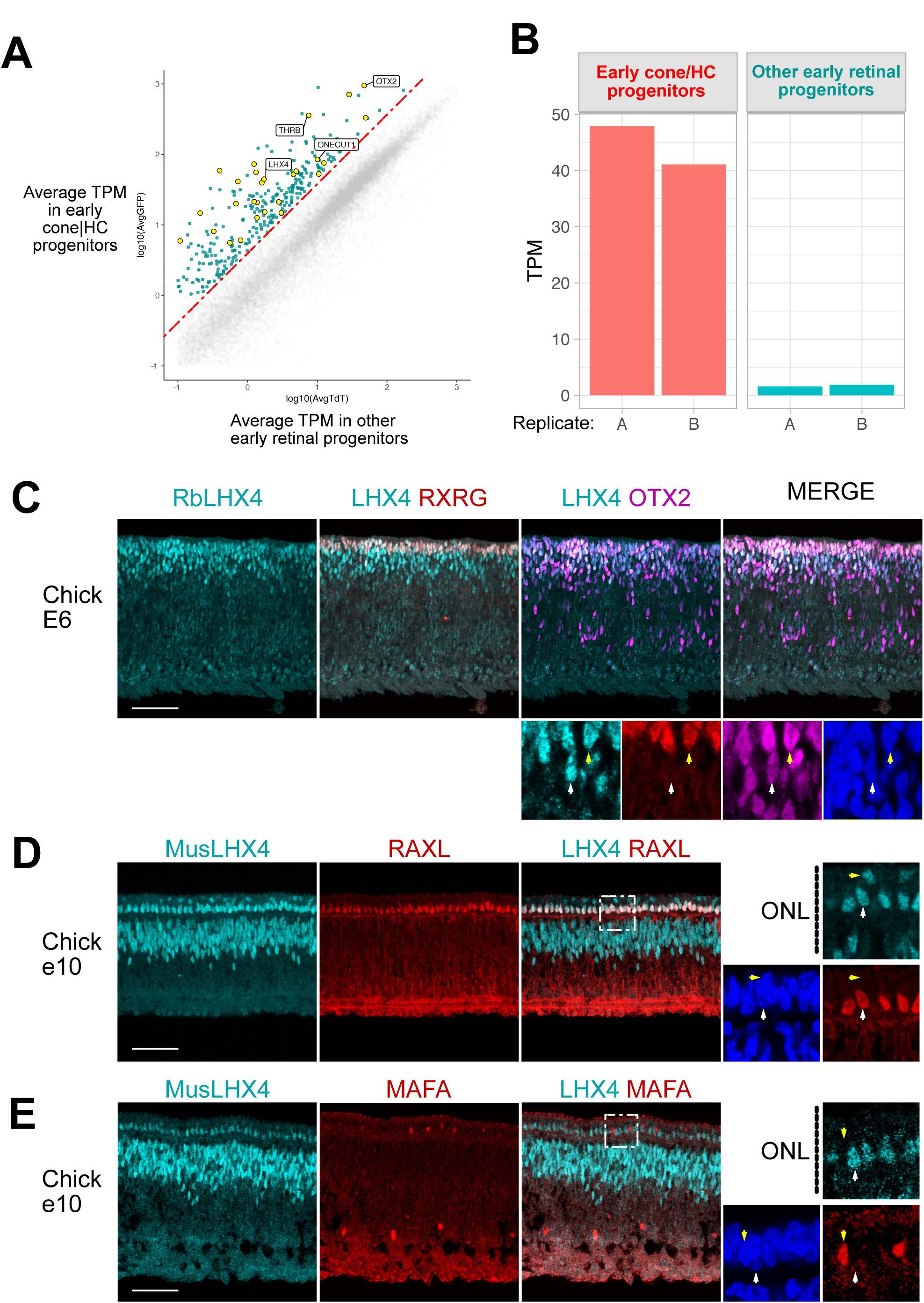
LHX4 is present in early cone photoreceptors during chick retinal development. Large panels are maximum intensity projections Z-stacks and small panels are single planes of the same Z-stacks (A) Average Transcripts Per Million (TPM) of genes between Cone/HC progenitors and other early retinal progenitors. Highlighted in green are genes 1.5-fold enriched in Cone/HC progenitors and highlighted in yellow, transcription factors. (B) TPM values for LHX4 in Cone/HC progenitors and other early retinal progenitors. (C) Cross-section of E6 chick retina imaged for LHX4, RXRG, and OTX2. Higher magnification panels show a single z-plane with marked a LHX4+/RAXL+ (white arrow) or LHX4+/OTX2+/RXRG+ cell (yellow arrow). (D) Cross-section of E10 chick retina imaged for LHX4 and RAXL. Higher magnification panels show a single z-plane with marked a LHX4+/RAXL+ (white arrow) or LHX4+/RAXL- cell (yellow arrow). (E) Cross-section of E10 chick retina imaged for LHX4 and MAFA. Higher magnification panels show a single z-plane with marked a
LHX4+/MAFA- (white arrow) or LHX4-/MAFA+ cell (yellow arrow). Scale bar represents 50 μm

A previous report has examined the presence of LIM-domain factors in early chick photoreceptor development^15^. This study suggested that LHX3 was abundantly present in the apical portion of the retina and localized to photoreceptors once the ONL is clearly distinguished. As the RNA-Seq data indicated that LHX4 expression is prominent in ThrbCRM1 reporter-positive cells at early stages while LHX3 transcript presence is marginal in all targeted cells (Supp. Fig. 1 A), we suspected that this previous study could have detected LHX4 instead of LHX3 at earlier timepoints. To test this, we electroporated a mouse LHX4 misexpression plasmid (CAG::mLHX4) plasmid alongside a CAG::nucβgal construct into the chick E6 retina, cultured it for 2 days, and detected for LHX4 using the LHX3 Developmental Studies Hybridoma Bank (DSHB) antibody. As predicted, we observed robust immunoreactivity of βgal+ electroporated cells with the LHX3 antibody (Supp. Fig. 1 B), suggesting that this antibody is also capable of detecting mouse LHX4, and thus likely chicken LHX4 as well.

With the use of this LHX3/4 antibody and a rabbit polyclonal antibody, we examined LHX4 presence during embryonic development of chick at E6 and E10. Expression at E6 is restricted to the scleral portion of the retina, where photoreceptors are located. As the LHX4 transcript was highly enriched in cells that activate the ThrbCRM1 element, which requires OTX2 for activity, we expected LHX4 to be present predominantly in the OTX2+ population at E6. Indeed, LHX4 is detected in a large percentage (but not all) of OTX2-positive cells (Fig. 1 C).

At E6, cones are the major class of photoreceptors that are produced, as the earliest known rod photoreceptor marker in the chicken retina, L-Maf (MAFA), is not detected until E9^16^. As the LHX4 pattern at this stage is localized in the apical portion of the retina, we examined LHX4+ cells for co-expression with the cone marker RXRG. Many of the most apically located LHX4 cells were indeed positive for RXRG (Fig. 1 C).

It is unclear to what extent and in which cells LHX3, LHX4 or both proteins are present in later timepoints when the transcriptional status of LHX3 may change. In one report^15^, antibody staining using DSHB anti-LHX3 showed strong nuclear Outer Nuclear Layer (ONL) and INL signal at E10, which suggested photoreceptor and bipolar signals. We also detected signal in the ONL and INL with the LHX3 antibody. However, the rabbit anti-LHX4 antibody detected a similar pattern with strikingly strong signal in the ONL and weaker in the INL (Suppl. Fig. 1 C). As clear evidence of LHX3 RNA expression in the ONL was not observed in the previous study, this suggests that LHX4 and not LHX3 may continue to be expressed in E10 chicken photoreceptors.

To examine the photoreceptor subtype expression more closely, we compared to RAXL expression, a marker for cone photoreceptors^16^, and observed that all RAXL-positive cells were also positive for LHX4 (Fig 1 D). A smaller number of cells in the upper part of the ONL were LHX4-positive and not RAXL-positive. As the ONL contains both cones and rods, we used an antibody to MAFA^16, 17^, the earliest rod marker, to determine if these cells were rod photoreceptors. No overlap between MAFA and LHX4 was detected at E10 in the ONL (Fig 1 E). This data suggests that LHX4 is expressed predominantly in developing cone photoreceptors in the chicken retina.

### LHX4 is present in early cone photoreceptors in the mouse retina

We sought to establish if this protein was also present in early photoreceptors of the mouse retina. At E14.5, LHX4 immunoreactivity was present in the scleral portion of the retina, where developing photoreceptors are located (Fig 2 A). To confirm that LHX4 was present in cone photoreceptors, we used the cone-expressed genes RXRG and OTX2 to identify these cells^18^. RXRG is also expressed in some retinal ganglion cells but these cell types can be readily distinguished by location. Many LHX4 cells are positive for RXRG and OTX2 at E14.5, in a similar pattern to the chick retina. At E14.5, we found that 91.1+-1.9% (mean+-SEM) of cells positive for LHX4 were also positive for RXRG (Fig 2 B), signifying that the majority of the LHX4+ population at this timepoint were composed of early cones. In fact, nearly all cells positive for RXRG at E14.5 were also positive for LHX4 (99+-1%, Fig 2 C).

**Figure 2.**
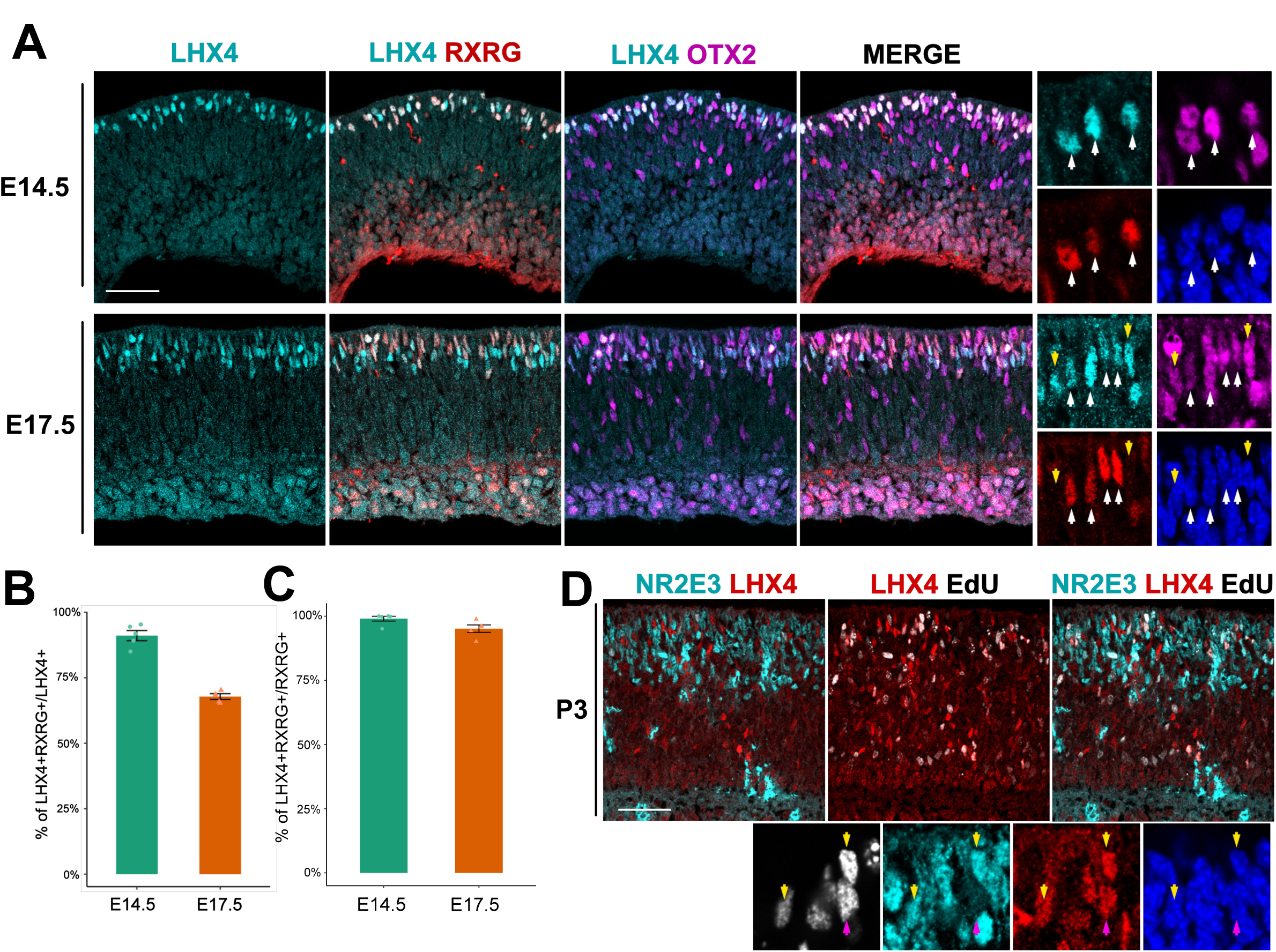
LHX4 protein is present in early photoreceptors precursors during mouse retinal development. Large panels are maximum intensity projections Z-stacks and small panels are single planes of the same Z-stacks A. Cross-section of embryonic mouse retinas imaged for LHX4, RXRG, and OTX2 at the designated timepoints. Higher magnification panels show a single z-plane with marked LHX4 cells positive (white arrows) or negative (yellow arrows) for RXRG and OTX2. B. Quantification of the percentage of LHX4+ cells that are also RXRG+ at E14.5 and E17.5. C. Quantification of the percentage of RXRG+ cells that are also LHX4+ at E14.5 and E17.5. D. Cross-section of a P3 retina after a P0 EdU injection imaged for EdU, NR2E3 and LHX4. Higher magnification panels show a single z-plane with marked LHX4+/EdU+ cells positive (yellow arrows) or negative (purple arrows) for NR2E3. Scale bar represents 50 μm

At a later embryonic stage (E17.5) we found that LHX4 cells still co-localize with RXRG protein, but there was an increase in LHX4+/OTX2+ cells that did not express RXRG (Fig 2 A). At this timepoint, only 67.8+-1.1% of LHX4+ cells were positive for RXRG, suggesting that LHX4 was active in other emerging cell populations in addition to early cones (Fig 2 B). However, 95.1+-1% of RXRG+ cells were positive for LHX4 at e17.5, indicating that the majority of cones retained LHX4 expression (Fig 2 C). The rabbit LHX4 polyclonal antibody had a reduced quality of staining at this timepoint, so it is possible the small fraction of cones that are not LHX4+ had undetectable LHX4 signal under these conditions or could point to a small subpopulation of cones that do not express LHX4. Additionally, we tested if LHX4 was present in post-mitotic cone photoreceptors or in dividing cells. E14.5 retinas exposed to a 2 hour 5-ethyny-2’- deoxyuridine (EdU) pulse did not show a qualitative overlap between EdU and LHX4 or RXRG (Supp. Fig. 2), which is consistent with LHX4 expression beginning after cell cycle exit. We conclude that LHX4 is a relatively specific marker for post-mitotic cone photoreceptors at early stages in the mouse retina but is also present in other cell populations at later developmental stages.

### LHX4 is expressed in several developing and adult cell types

Previous reports indicated that LHX4 protein was present in adult cone BCs^19^, but little is known about its expression pattern in the developing mouse retina. We also observed strong LHX4 labeling in the upper portion of the INL in cells positive for OTX2, an adult marker for BCs (Supp. Fig. 3). Interestingly, we also noticed sparse, but positive staining in the ONL. This LHX4 signal was located in some but not all RXRG+ cones with varying degrees of strength (Supp. Fig. 3) suggesting that a sub-population of cones maintain LHX4 expression.

**Figure 3.**
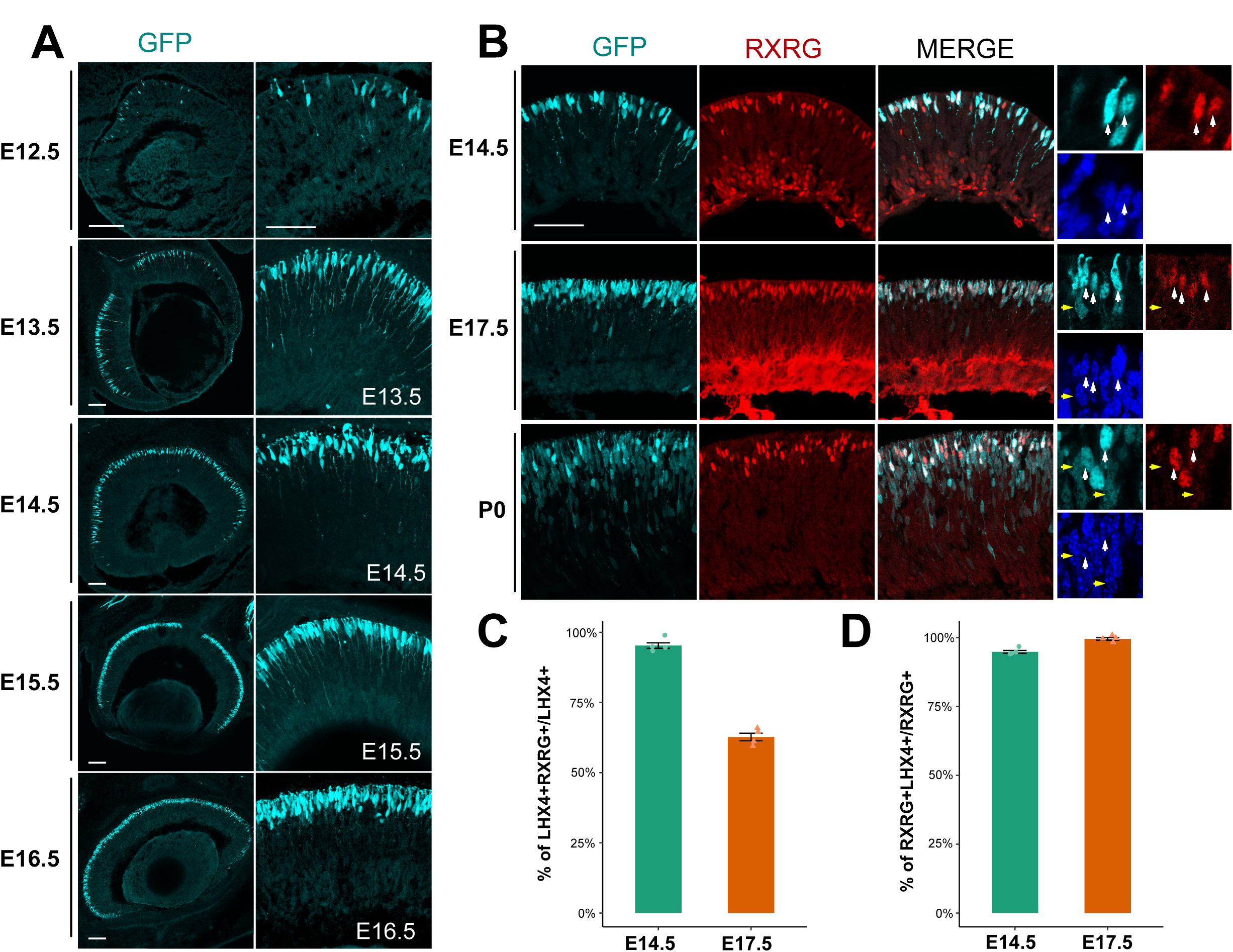
LHX4 BAC-EGFP is a reliable reporter of photoreceptors during retinal development. Large panels are maximum intensity projections Z-stacks and small panels are single planes of the same Z-stacks. Scale bar represents 50 μm A. Cross-section of embryonic LHX4-GFP mouse retinas imaged for EGFP at the designated timepoints. B. Cross-section of embryonic LHX4-GFP mouse retinas imaged for EGFP and RXRG at the designated timepoints. Higher magnification panels show a single z-plane with marked EGFP cells positive (white arrows) or negative (yellow arrows) for RXRG. C. Quantification of the percentage of EGFP+ cells that are also RXRG+ at E14.5 and E17.5. D. Quantification of the percentage of RXRG+ cells that are also EGFP+ E14.5 and E17.5.

As noted above, reports^15, 20^ suggested that LHX4+ cells in the INL are likely developing BCs. In the mouse, our data indicated that LHX4 is initially in developing cones. However, at E17.5, RXRG cones no longer represent the near totality of LHX4+ cells. Since the peak of BC production is not seen until ∼P3^21^, the identity of the remaining cells is unclear. While BCs can be produced at earlier timepoints (including E17.5), we still sought to ascertain if only LHX4+ BCs were being produced at this time, or if this LHX4 expression is present in another cell type. As rods are another OTX2+ cell type produced at this time, we aimed to see if these cells expressed LHX4 during development. Newborn mice (P0) were injected with EdU to mark cells undergoing S-phase at the time of injection and 3 days later the retinas were harvested. This length of time was chosen to allow some newly produced rods enough time to produce NR2E3, a well-known marker for rod fate^22^. It has been previously shown that NR2E3 is also present in cone photoreceptors transiently^23–25^, but, as determined previously ^26^, no cones are produced at P0. Therefore, EdU and NR2E3 co-localization should reliably mark cell types other than cones, likely rod photoreceptors. NR2E3 expression has not been reported in BCs but given the similarities in molecular profiles between BCs and photoreceptors, at this time we cannot exclude this possibility. We observed LHX4+ cells that are positive for NR2E3 and EdU at P0 (Fig 2 D). This indicates that while LHX4 is present in developing and adult BCs and cones, it is also possible that is transiently expressed in some rod photoreceptors.

### The LHX4::GFP transgenic line recapitulates the endogenous LHX4 expression pattern

We set out to establish if a reported LHX4::EGFP mouse transgenic line^14^ could be a reliable tracer for early developing cones. The LHX4::GFP line had robust expression in the retina throughout development (Fig 3 A) and GFP signal was located in the apical portion of the retina, where photoreceptors are located during embryonic development. EGFP could be detected during embryonic and postnatal development and also had stable expression in the adult retina (Supp. Fig. 4). This line has been used previously for transcriptomic and electrophysiological analyses^27, 28^, because it shows reliable activity in cone BCs. Interestingly, a subpopulation of cones is labeled by this line in the adult and this selective expression resembles the sparse presence of LHX4 protein in the adult mouse retina. To verify if this sparse labeling was due to LHX4 presence in exclusively S or L/M types of cones, which are located in contrasting gradients in the adult retina, we assessed EGFP expression in the dorsal and ventral retina^29^. We detected no difference in EGFP expression between these two areas and was present in a subset of Cone-Arrestin+ cells regardless of Dorsal-Ventral position (Supp. Fig 5).

**Figure 4.**
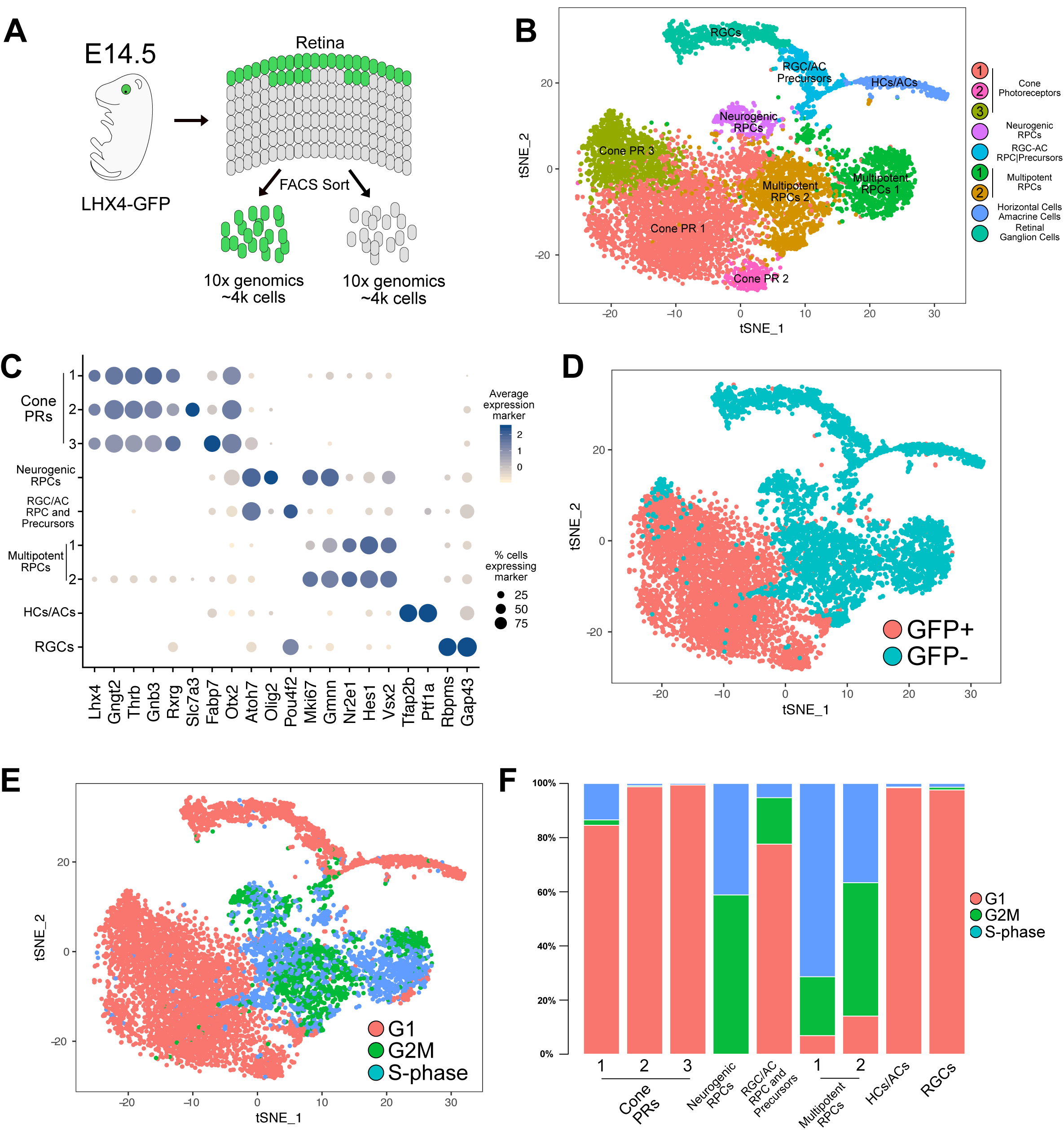
TSNE - Single cell sequencing of LHX4-GFP E14.5 developing retina. (A) Schematic of experimental design for single cell collection and sequencing. (B) tSNE plot of all GFP+ and GFP- cells collected, analyzed in tandem, and displaying the results of unsupervised cluster analysis with the assigned cell class according to their molecular signature. (C) Dot plot displaying the average expression and percentage of cells expressing specific markers in each cluster. Markers displayed on the x axis were used for assignment of cell class to each cluster. (D) Same tSNE plot as (B) displaying the original source of each cell, GFP+ or GFP- samples. (E) Same tSNE plot as (B) displaying the cell cycle assignment of each cell. (F) Percentage of cells in each cluster assigned to different cell cycle phases.

**Figure 5.**
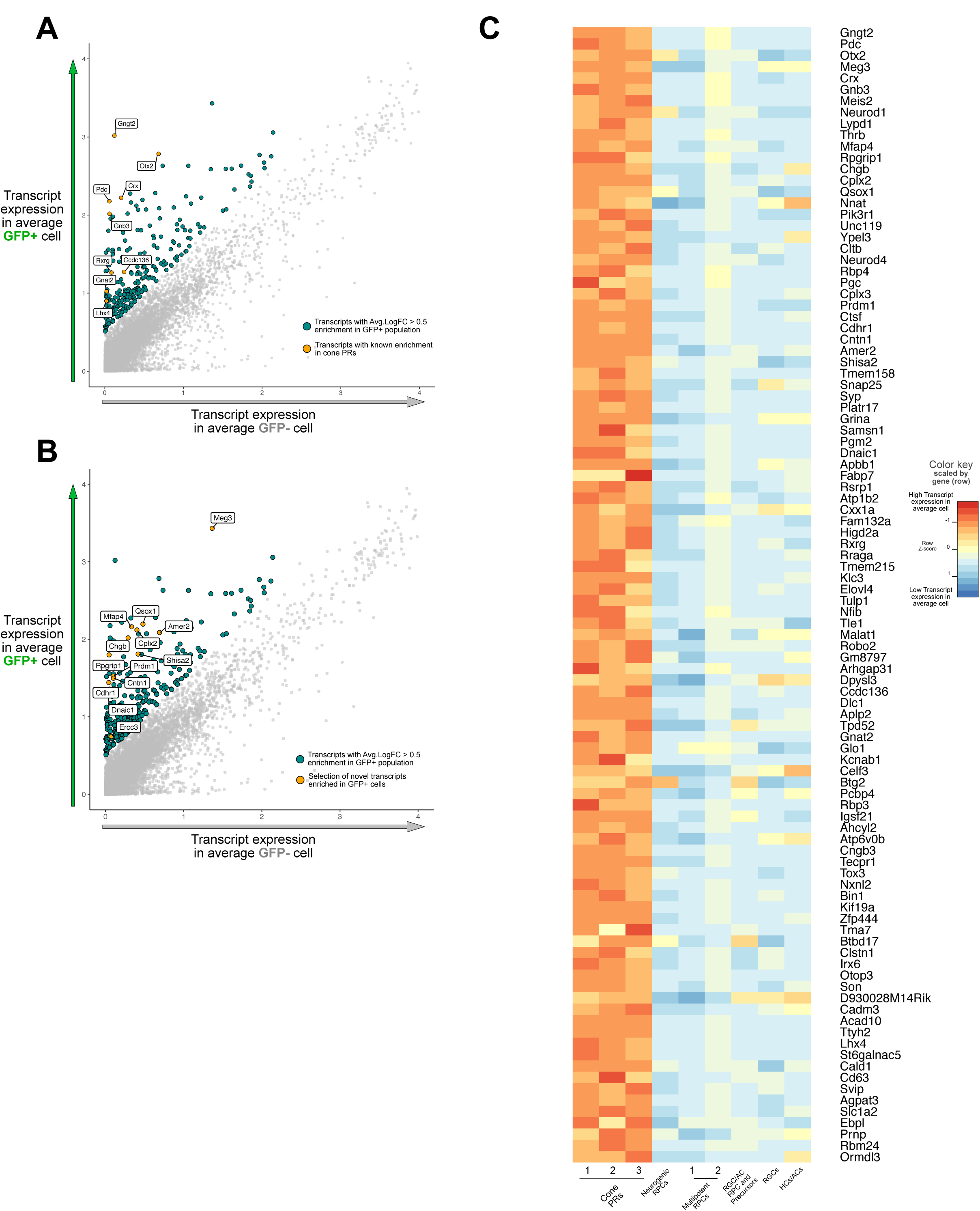
Identification of known and novel transcripts enriched in cone photoreceptors in LHX4-GFP+ cells. (A) Plot displaying the expected transcript expression in an average GFP+ (y-axis) or GFP- (x-axis) cell. Highlighted are all genes with >0.5 fold change enrichment in GFP+ cells and labelled are a selection of cone-related transcripts. (B) Same plot as (A) labeling novel transcripts enriched in the same population. (C) Heatmap of top 100 transcripts enriched in GFP+ cells. Color indicates the expected transcript expression of that gene in an average cell for each cluster identified. Expression values are scaled per gene.

We were unable to use the rabbit anti-LHX4 antibody to validate the LHX4::GFP expression as we encountered abnormal LHX4 immunoreactivity; not nuclear, as expected, but cytoplasmic and precisely overlapped with EGFP expression. Noted in the strain description, EGFP was placed after the ATG of LHX4, replacing the LHX4 coding region so as to function as a tracer. We amplified and sequenced the coding region downstream of the LHX4 ATG using the reported 5’ UTR primer for genotyping and, surprisingly, found it contained a 70 base pair section of 5’ endogenous LHX4 coding sequence immediately followed by a multiple cloning site and the EGFP coding sequence, all in frame to produce a LHX4::GFP fusion protein. Thus, the first 34 amino acids of this fusion protein are from the sequence of LHX4 and the multiple cloning site (Supp. Fig. 6 A).

To test if this portion of LHX4 contained an epitope for the LHX4 antibody, we amplified the coding sequence from the LHX4::GFP genomic DNA, cloned it into a misexpression vector, and electroporated it into the chick retina (Supp. Fig. 6 B-D). Control retinas had no detectable EGFP and normal staining of LHX4 with the rabbit antibody (Supp. Fig. 6 B). In contrast, cells electroporated with the CAG::5pLHX4GFP construct were strongly immunoreactive to the LHX4 antibody in a pattern that completely overlapped that of the cytoplasmic EGFP signal (Supp. Fig. 6 C). The rabbit- LHX4 still detected endogenous LHX4, as we imaged the same retinas at the edge of the electroporation patch with higher gain and observed endogenous protein expression alongside overexposed electroporation signal (Supp. Fig. 6 D).

Therefore, to assess if EGFP recapitulates LHX4 expression patterns and its activity in early cones, we resorted to examination of RXRG expression in EGFP+ cells at two relevant timepoints of embryonic development. At E14.5, the EGFP reporter faithfully recapitulated LHX4 expression as 95.3+-1% of all EGFP+ cells were positive for RXRG (Fig 3 C). Likewise, at E17.5 only 65.7+-1.3% of EGFP cells were RXRG +. As the LHX4 immunodetection showed, nearly all RXRG+ cones are positive for LHX4 at both E14.5 and E17.5. This was also true in the LHX4::EGFP+ population, where 94.8+-0.5% and 99.5+-0.5% of all RXRG cells, respectively, were positive for the EGFP reporter (Fig 3 D). Taken together, this data suggests that the LHX4 BAC-EGFP reliably recapitulates LHX4 protein expression and is a dependable marker for cone photoreceptors during early retinal development.

### Single cell sequencing of LHX4::GFP cells in the E14.5 developing mouse retina

Having verified that the LHX4::GFP reporter is a marker for cone photoreceptors in the early stages of mouse retinal development, we took advantage of this system to examine the molecular profile of early cone photoreceptors using single cell RNA sequencing. LHX4::GFP E14.5 littermates were screened for GFP expression and positive retinas were pooled and dissociated in preparation for FACS sorting (Fig 4 A). GFP+ and GFP-cells were collected and approximately 4000 cells were sequenced per condition using the 10x Chromium platform.

Using the Seurat program^30^, we performed an unsupervised clustering analysis on the combined cell transcriptomes from both conditions that passed standard 10x QC (GFP+: 3728, GFP-:4444). TSNE projections revealed that the GFP+ and GFP-populations clearly segregated, consistent with LHX4::GFP cells being a molecularly distinct population (Fig 4 B). The analysis separated the cells into 9 distinct clusters, which were identified by established markers and bore hallmarks of known cell classes in the developing retina (Fig 4 C-D; Supplementary File 1).

Two cell clusters were assigned as multipotent RPCs based on expression of multipotent RPC markers such as VSX2, HES1, NR2E1, among others, as well as a large percentage of cycling cells ^31^ (Fig 4 E-F). One other cluster was predominantly comprised of cycling cells, with markers such as OLIG2, NEUROG2, OTX2 and ATOH7, which identify these as likely neurogenic RPCs^32, 33^ with limited mitotic potential. A second cluster had a smaller fraction of cycling cells, high ATOH7 levels and expression of POU4F2, but no OTX2 or OLIG2, which we assigned to RGC/AC RPCs and Precursors as they exhibited markers of likely differentiation to RGCs or ACs.

Of the cell clusters assigned as mostly post-mitotic, one was assigned as RGCs as it displayed known markers RBPMS and GAP43^34, 35^. Another assigned to AC/HCs, exhibiting markers for both these fates, like TFAP2B and PTF1A^36, 37^. As previously reported^33^, while ACs and HCs are distinct fates, it is difficult to separate them by their transcriptomes in early development and without an appropriate amount of sequenced cells for proper resolution.

Three clusters were assigned as cone photoreceptors. As expected from our previous data, these consisted of the near totality of GFP+ cells and displayed markers of cone photoreceptors: THRB, RXRG, GNGT2 and GNB3 (Fig 4 C-D), as well as LHX4. We did detect some sparse Nrl-positive cells within this population, possibly reflecting some activity of the LHX4 reporter in rod photoreceptors. The cell cycle phase of these clusters is consistent with our previous data suggesting LHX4 is in post-mitotic cones at E14.5. As a result of the sorting strategy, cone representation was high which allowed the clustering analysis to resolve cone subpopulations. All cone clusters had high levels of established markers, but two subpopulations differed in expression of genes. The highest differential marker for one of the populations was FABP7, a previously characterized marker for cones in the adult murine retina^38^ with reported expression in the developing retina^39^ but not at this early stage. Meanwhile, a second subpopulation had increased levels of solute-carrier genes like SLC7A3 and SLC7A5 (Fig 4 C). Our analysis indicates that we successfully sorted and sequenced a developing E14.5 retina and identified its cell populations while enriching for cone photoreceptors.

### Early cone marker identification in LHX4::GFP cells

We sought to use this dataset to find new markers for early cone photoreceptors. Using Seurat, we performed a differential expression analysis, comparing the GFP+ population, which we established as cones, with the GFP-population, which should encompass the other concurrent populations in the developing retina. Additionally, for visualization purposes and as a proxy for average population levels of transcript expression we calculated the expression of an average GFP+ and a GFP-cell and used this in combination with the differential expression results. We identified over 898 significantly differentially expressed transcripts with enrichment in cones. As expected, we identified known cone markers as highly enriched in the GFP+ population (Fig 5 A, Supplementary File 2). We then looked for novel transcripts enriched in this population (Fig 5 B). A heatmap for the top 100 cone-enriched transcripts with the transcript expression of an average cell in every individual cluster is included in Fig. 5 C.

The above analysis identified genes enriched in early cone photoreceptors compared to the other cell types present at that time. We first compared these results to other studies that have reported cone transcriptome analyses in the mouse or human (Supp Fig 7 A-B). Only one study has reported a transcriptome from early developing cones, similar to this one^4^, through the use of human fetal explants infected with cone opsin promoter reporter viruses, determining early and late phases of gene expression in labeled versus non-labeled cells. A number of genes are enriched in both datasets (165 genes in early and 211 in late fetal cones), suggesting that the mouse is a potential model for investigating the function of these genes in cone photoreceptors. (Supp. Fig 7, Supp. File 3 and 4).

Additional datasets of purified adult rods and cones from the mouse have been used to identify differential transcripts between these two types of photoreceptors^7^. Comparison to this dataset identified that 251 of the genes in our dataset were enriched in adult cones and 121 were enriched in adult rods (Supp. Fig 7, Supp. File 3 and 5). Adult cone-enriched genes present in this dataset support the cell-specificity of known early markers detected in our dataset, such as RXRG, as well as many genes not explored in detail before, such as QSOX1 or LHX4.

We next were interested in determining whether the genes associated with early cone genesis were negatively regulated by the rod photoreceptor factor Nrl. A prevalent model for Nrl function is that it serves as a fate switch in photoreceptor precursors, with Nrl-negative precursors becoming cones and Nrl-positive ones becoming rods^8, 12, 40^. In Nrl mutants, the photoreceptors that normally would become rods are found to undergo a morphological and gene expression change that could suggest a rod to cone fate switch. However, it has been noted that these cells are morphologically distinguishable from normal cones and, in addition, known early cone genes, such as Thrb and Rxrg, are unaltered in newborn photoreceptors, which suggests that Nrl may not be a master regulator of this fate choice ^8, 12, 41^. As there are few known early cone genes, we sought to identify other cone photoreceptor genes, in addition to Thrb and Rxrg, that are regulated independently of Nrl. We first identified those genes in our dataset that were dysregulated either positively or negatively in Nrl knockout photoreceptors. The Nrl dataset used contained multiple isoforms of genes, but to apply a stringent criterion, any isoform that showed dysregulation at either P2 or P28 in Nrl mutants led to that gene being removed, leaving a total of 259 Nrl-independent genes (Fig 6 A). Interestingly, many of the cone-enriched genes present at E14.5 are only altered in Nrl mutants at P28 and not also at P2 when a large number of rods would be generated (see discussion).

**Figure 6.**
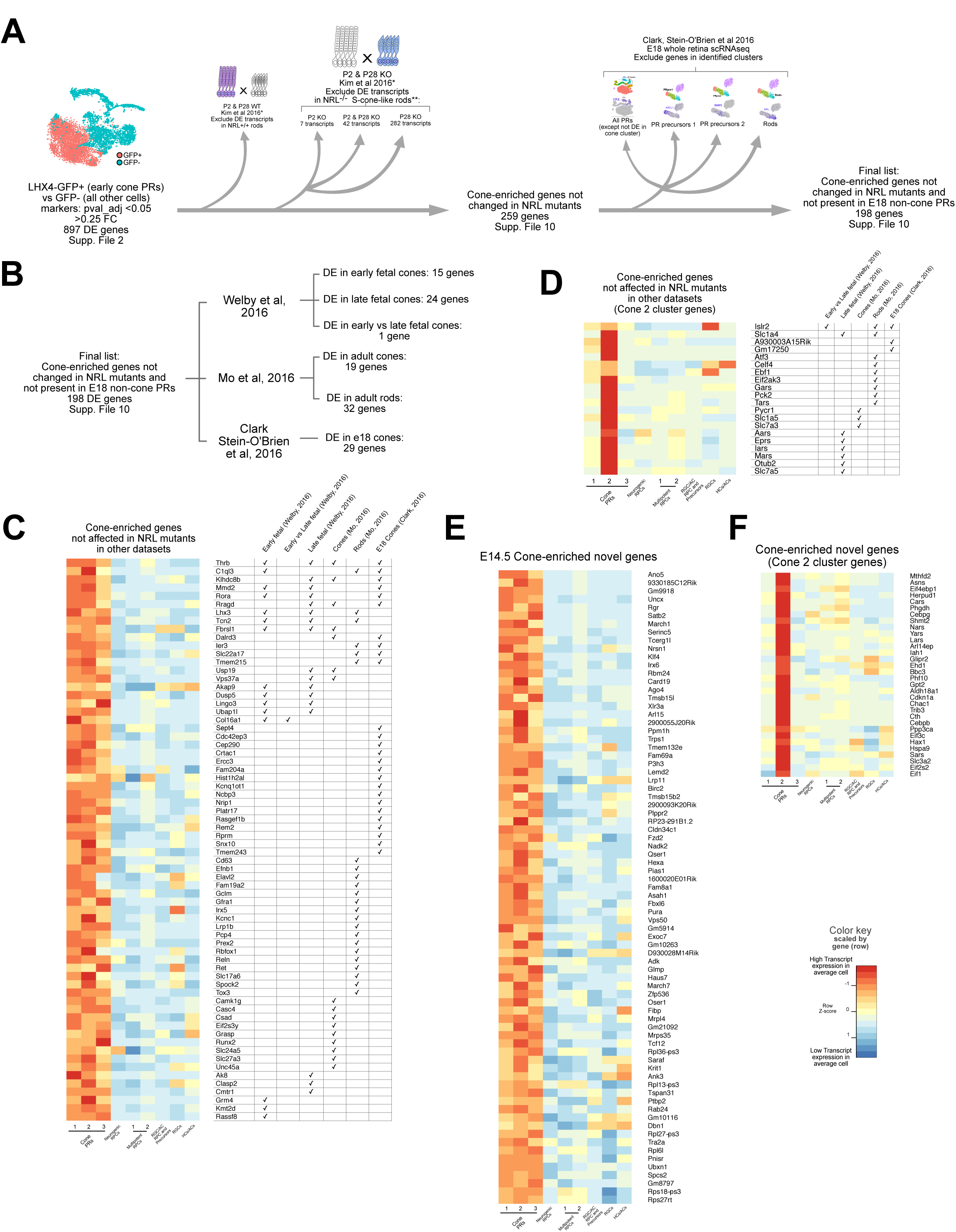
E14.5 genes not changed in NRL mutants and novel cone-enriched genes. (A) Schematic describing the comparison and selection of genes enriched in E14.5 cones that are not changed in the NRL mutant dataset and not present in early rods or precursors at E18 (Clark, Stein O’Brien 2018) (B) Amount of genes found overlapping in the displayed datasets from the final list of genes. (C) Heatmap of transcripts enriched in E14.5 cones that are not changed in the NRL mutant dataset and not present in early rods or photoreceptor precursors at E18. The yed are also present in the datasets analyzed, and their presence is indicated by a checkmark in the right table. Color indicates the expected transcript f that gene in an average cell for each cluster identified. Expression values are scaled per gene. (D) Same as C but for genes found in the Cone 2 cluster. (E) Heatmap of novel genes found in this study to be enriched in E14.5 cones but not present in any of the other datasets. Color indicates the expected transcript f that gene in an average cell for each cluster identified. Expression values are scaled per gene. (F) Same as E but for novel genes in the Cone 2 cluster.

As some of these genes likely represent pan-photoreceptor genes that wouldn’t be expected to be under the transcriptional control of Nrl, we identified those genes also expressed in developing rods. As an identifiable rod cluster was not present in our E14.5 dataset, we analyzed a recently generated single cell dataset from E18.5 whole retinas^33^. The data from 2 replicates of E18 was extracted and we performed an unbiased clustering analysis using Seurat^30^. (Supp Fig 8). The detected 9 clusters were assigned to known cell types through marker expression. Within those, two clusters were positive for CRX+ and, thus, determined to be likely photoreceptors. First, all DE transcripts for photoreceptors at e18 were determined (Supp. Table 7, Supp. Fig 8) and then these cells were re-analyzed for further sub-clustering. We were able to determine a cluster with cone signature, rod signature and 2 precursor clusters. From those, we determined all DE transcripts between photoreceptor clusters (Supp. Table 8, Supp Fig 8). Interestingly, many of the genes detected as enriched in early cone photoreceptors in this study have been identified as differentially expressed across retinal pseudotime by Clark, Stein O’Brien et al, 2018 (Supp. Table 9)

Removal of genes with expression in any of the non-cone clusters at e18, produced 198 Nrl-independent cone-enriched genes (Fig 6 A-B, Supp. Table 10). To provide some measure of confirmation of the cone/photoreceptor-associated expression of these genes, we identified those genes with cone-enriched expression or specifically enriched in cone cluster 2 that were also identified in the previously mentioned datasets (Fig 6 C, D, Supp. Table 10). This left a set of genes with cone-enriched expression or specifically enriched in cone cluster 2 that have only been identified to date in this study as cone-associated (Fig 6 E, F).

## DISCUSSION

Our current knowledge of the molecular and cellular events involved in cone photoreceptor development is incomplete. One missing dataset has been a comprehensive gene expression analysis of endogenously developing cones. Without such knowledge, the rational design of strategies to induce cone formation and adequate benchmarks to assess these cones, such as from stem cell cultures, are not possible. Here we identify LHX4 as a novel and highly specific marker of cone photoreceptors in the early stages of their development and the LHX4::EGFP reporter mouse as a tool to detect and analyze these cells in the developing mouse retina.

The single cell transcriptome analysis allowed for an unprecedented molecular examination of early cone photoreceptors. With the increased representation that the LHX4::GFP mouse permitted for the purification of this relatively rare cell type, we were able to identify sub-clusters of cones at this timepoint. The biological significance of these sub-clusters, however, remains, to be determined. They could reflect temporal differences in the differentiation process or spatial effects linked to the dorsal-ventral position of cells, which is known to influence cone opsin expression at later timepoints. One distinct cluster upregulates the SLC7A3 and SLC7A5 genes. SLC7A5 is a known thyroid hormone transporter for T3/T4 states^42^ that has been reported in human fetal retinas and organoids^43, 44^. Because these transcriptomic analyses were done in whole retinas it wasn’t clear if this expression was specific to any particular cell population, but our data suggests that mouse cone photoreceptors specifically express these genes. SLC7A3 has not been reported in the retina but likely plays a similar role as its expression has been linked to T3 administration in the brain^45^. Thyroid hormone and its targets are known to affect M-cone differentiation^43, 46–49^. We found several other thyroid-related genes throughout the retina, but only THRB was exclusive to cones at this time. For example, there was almost no detectable amount of DIO2 in any cell but many more counts of DIO3, although these were present in the multipotent RPC clusters. Another observation of cone heterogeneity was in the expression of LHX4 protein and the LHX4::GFP reporter in more mature retinal tissue. Whether this sustained expression of LHX4 in subsets of cones is biologically relevant or is an epigenetic or temporal phenomenon without a functional significance warrants further study.

There is a major interest in understanding the molecular differences between cone and rod photoreceptors and how these differences arise during development. This would provide insights into how these two classes of photoreceptors evolved as well as inform the strategic development of methods to specifically produce these cell types. The predominant photoreceptor determination framework is that cones and rods developmentally diverge based solely on whether newborn photoreceptors begin to express the Nrl transcription factor or not. The analysis performed here with multiple previously published datasets and the current one suggests that endogenous newborn cones have a unique molecular signature compared to the induced cones that form in the Nrl knockout. There are caveats that must be considered. For one, the current study used a massive enrichment of cones, which allowed for increased power to detect gene expression differences between cones and other cells. Comparison to other datasets that had less statistical power may have led to the false conclusion that a gene was not differentially expressed. In addition, we compared the profile of embryonic cones to Nrl-dysregulated genes at P2. There could be currently unidentified signaling factors that are temporally different and influence gene expression that could account for these differences. It is important to note that a large number of cones genes are in fact dysregulated in the Nrl mutant. However, it is interesting that many of these genes found in early cones, such as RXRG, are not dysregulated in early rods, but are only significantly changed in much more mature rods. What the cause of this delay is and whether it impacts the differentiation of these transformed cells to a more bona fide cone photoreceptor signature is not known. Regardless, these differences warrant further examination of the utility and validity of using photoreceptors from the Nrl knockout model as a substitute for naturally formed cones.

## Materials & Methods

### Animals

All experimental procedures were carried out in accordance with the City College of New York, CUNY animal care protocols under IACUC protocol 965. CD-1 mice were used and provided by Charles River. The LHX4::GFP strain is a Bacterial Artificial Chromosome (BAC) insertion and (strain: Tg(Lhx4-EGFP)KN199Gsat/Mmucd, RRID:MMRRC_030699-UCD) was obtained from MMRRC. LHX4::GFP mice were kept and used experimentally only as heterozygotes. Fertilized chick eggs were from Charles River, stored in a 16°C room for 0-10 days and incubated in a 38°C humidified incubator. All experiments that used animals were not sex-biased.

### Genotyping

Genotyping was performed as specified in the MMRRC strain description page referenced (LHX4::GFP strain: Tg(Lhx4-EGFP)KN199Gsat/Mmucd, RRID:MMRRC_030699-UCD).

### Cloning and DNA electroporation

Misexpression plasmids were created by PCR amplifying the coding sequence for each gene with added AgeI/NheI or EcoRI sites. They were subsequently cloned into a CAG backbone, from which the GFP had been removed from Stagia3^47, 50^ and the CAG promoter had been cloned upstream. For mouse LHX4 the primers were designed against the annotated mRNAs in NCBI galgal5 RNA libraries with an added Kozak sequence 5’ (ACC). For 5pLHX4GFP, the 5’ primer from genotyping for LHX4::GFP mice was combined with a 3’ primer to the coding region of GFP and the fusion gene was amplified from genomic DNA extracted from LHX4::GFP+ mice. CAG::nucβgal was developed by the Cepko lab (Harvard Medical School). To deliver the plasmids after retinal dissection, ex vivo electroporation experiments were carried out as detailed in Emerson and Cepko, 2011^50^.

### Retinal cells dissociations and Florescence Activated Flow Sorting (FACS)

Dissociation of retinal tissue and FACS were performed as in Buenaventura et al, 2018^3^

### Immunohistochemistry and EdU labeling

All tissue processing and immunofluorescence experiments were performed as in Emerson and Cepko, 2011 and Buenaventura et al, 2018. For E14.5 mouse retinas, EdU labeling was performed by injecting pregnant dams with 150ul of 10mM EdU resuspended in 1X PBS. EdU detection was performed with a Click-iT EdU Alexa Fluor 647 imaging kit (C10340, Invitrogen).

### Microscopy

Confocal images were acquired using a Zeiss LSM880 confocal microscope using ZEN Black 2015 2.1 SP2 software and images were converted into picture format using the FIJI version of ImageJ^51^. Figures were assembled using Affinity Designer vector editor. Images were adjusted uniformly with regards to brightness and contrast.

### 10X single cell sample processing

Live cells were collected after dissociation and FACS sorting for GFP signal^3^. Samples were extracted from 2 litters of E14.5 LHX4::GFP mice, pooled into one sorting solution during dissociation. For each sample (GFP+ and GFP-), approx. 4000 cells were individually lysed, and their RNA transcribed and sequenced by the Columbia University Single Cell Analysis Core. Filtered count matrixes provided by 10X Cell Ranger pipeline using mm10 genome were used for downstream analysis.

### Single-cell clustering analysis

Unsupervised clustering analysis was performed using Seurat^30, 52^. Single cell transcriptomes from both LHX4::GFP E14.5 samples were analyzed in concert to produce cluster and TSNE analysis. Only cells with over 200 genes detected and only genes detected in >3 cells were used for initial loading of the cell matrices. Cell cycle was scored and was regressed out as an unwanted source of variation according to Seurat’s guidelines ^31^. For the LHX4-GFP dataset, cells were filtered for max. 3500 and min. 250 number of genes, and <0.25 percent mitochondrial content following standard guidelines for QC. nUMI, nGene were regressed out in addition to cell cycle. Using the established jackstraw procedure^53^ in Seurat, 30 PCs were chosen for clustering and TSNE projections. The ‘FindMarkers’ function was used with default parameters, only for positive markers, with the populations compared as described in the results section. For Clark, Stein O’Brien et al, 2018, similar procedures were used with the following changes: for the full dataset - 250 < nGenes < 3500, 0.075 < percent mito, 39 PCs and replicate effects were regressed out; for the photoreceptor subclustering – same as full but 6 PCs, data was extracted post-analysis of the full dataset.

### Data comparisons

For the comparison of retinal datasets, differential expression output data was obtained from each report^4, 7, 54^. The data was cross-referenced to the genes obtained in the analysis from this paper and are detailed in Supp. Files 3, 4, 5 and 6. In the case of Kim et al, 2016, genes are reported with PPDE as opposed to p-val-FDR or q-val as probability of differential expression, and were filtered to show only those with PPDE > 0.95

### Data availability

The data will be deposited in NCBI’s Gene Expression Omnibus^55^ and will be accessible through GEO Series accession number GSExxxxxx.

### Quantitative analysis of markers in retina sections

One z-stack from the central portion of each retina of four biological replicates was counted and used for mean and SEM calculations in each condition. Cells were counted using Cell Counter plugin in ImageJ or Fiji^51, 56^.

## Acknowledgments

Support was provided by NIH National Eye Institute grant R01EY024982 (to M.E.) and 3G12MD007603-30S2 (CCNY). The content is solely the responsibility of the authors and does not necessarily represent the official views of the National Eye Institute, the National Institute On Minority Health and Health Disparities or the National Institutes of Health. Excellent technical support was provided by Brandon Webley, Jeffrey Walker, and Jorge Morales. The MafA antibody was kindly provided by Celio Pouponnot. We thank Brian Clark and Seth Blackshaw for granting access to their recently generated single cell datasets.

