## Supplemental Figures for "Identification of genes with enriched expression in early developing mouse cone photoreceptors"

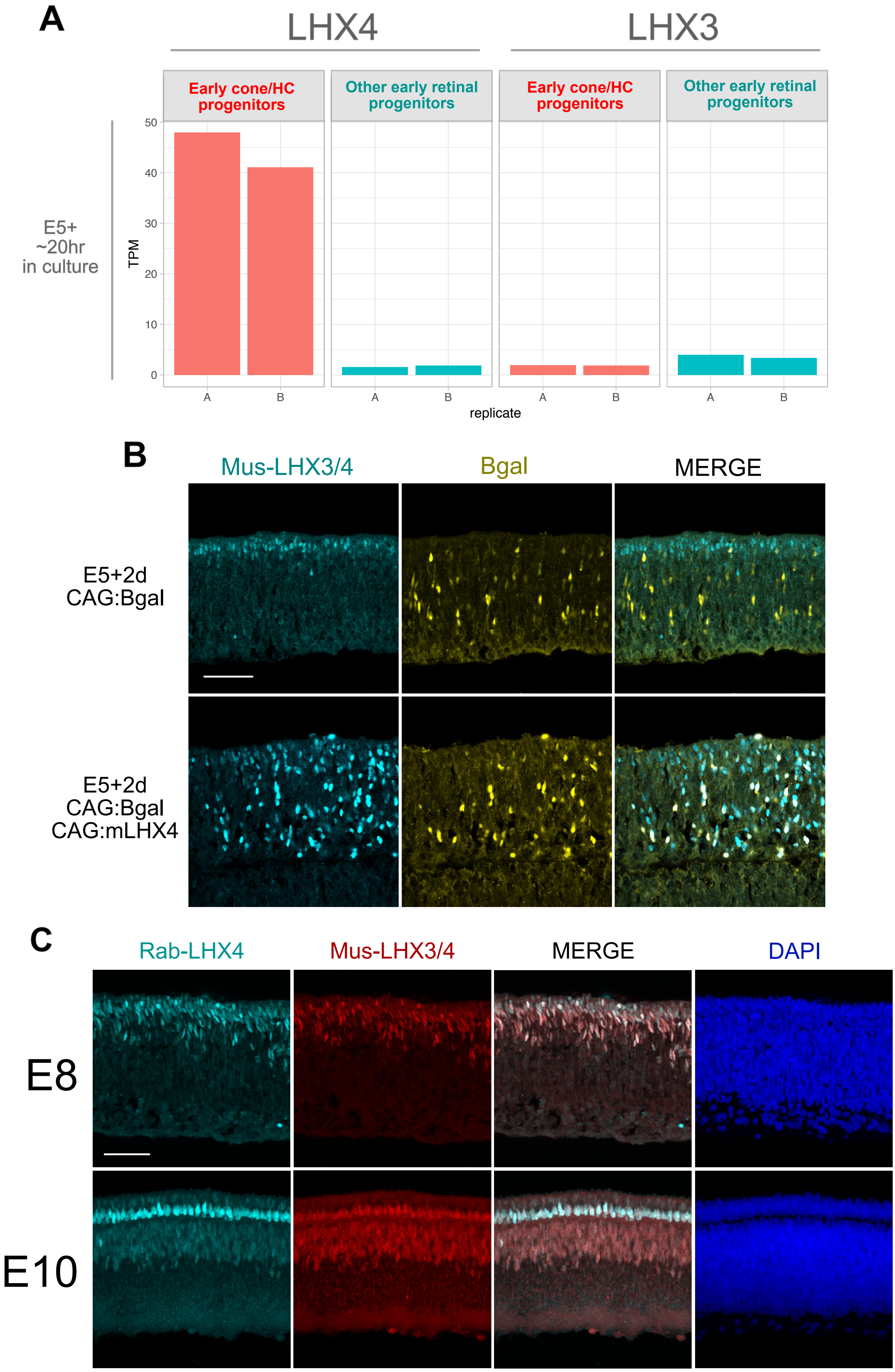


Supplemental Figure 1 - **LHX4 and not LHX3 is detected in early chick retina development.**

(A) TPM values for LHX4 and LHX3 in Cone/HC progenitors and other early retinal progenitors.

(B) Cross-section of a chicken retina electroporated at E5 and cultured for ~20hrs. Electroporated constructs are denoted on the left side. Retinas were imaged for Bgal and LHX3/4 with mouse antiLHX3 (DSHB).

(C) Maximum intensity projections of cross-section in chicken retinas at designated timepoints imaged for LHX4 with mouse (DSHB) and rabbit (Proteintech) antibodies.S cale bar represents 50 μm

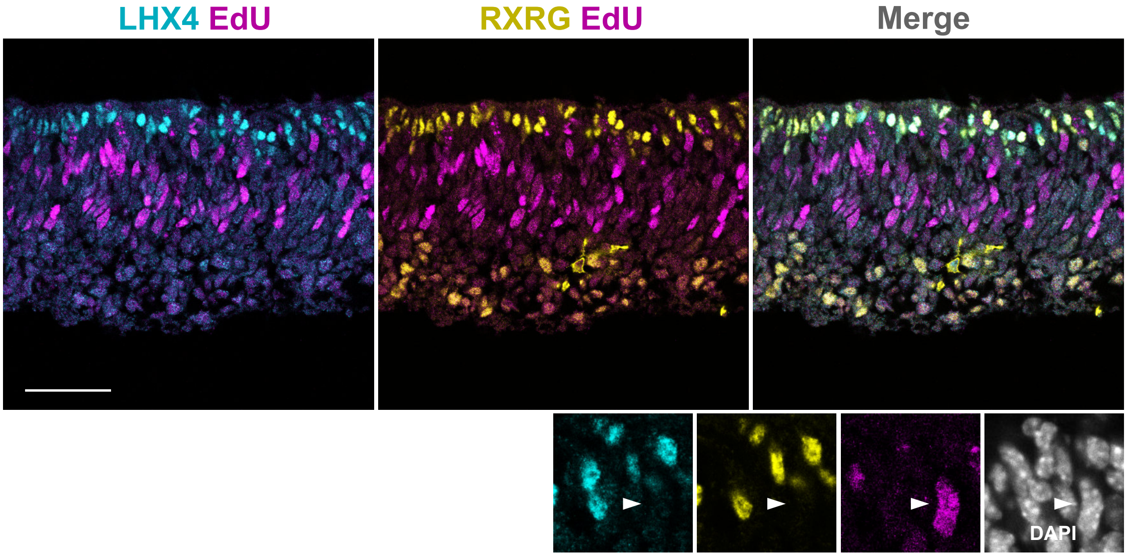


Supplemental Figure 2 - **LHX4 expression starts in post-mitotic cones**

Cross-section of a E14.5 mouse retina, developed for EdU (2hr pulse) and imaged for LHX4 and RXRG. Small panels are digitally zoomed. Arrow points at representative EdU cell negative for LHX4 and RXRG Scale bar represents 50 µm .

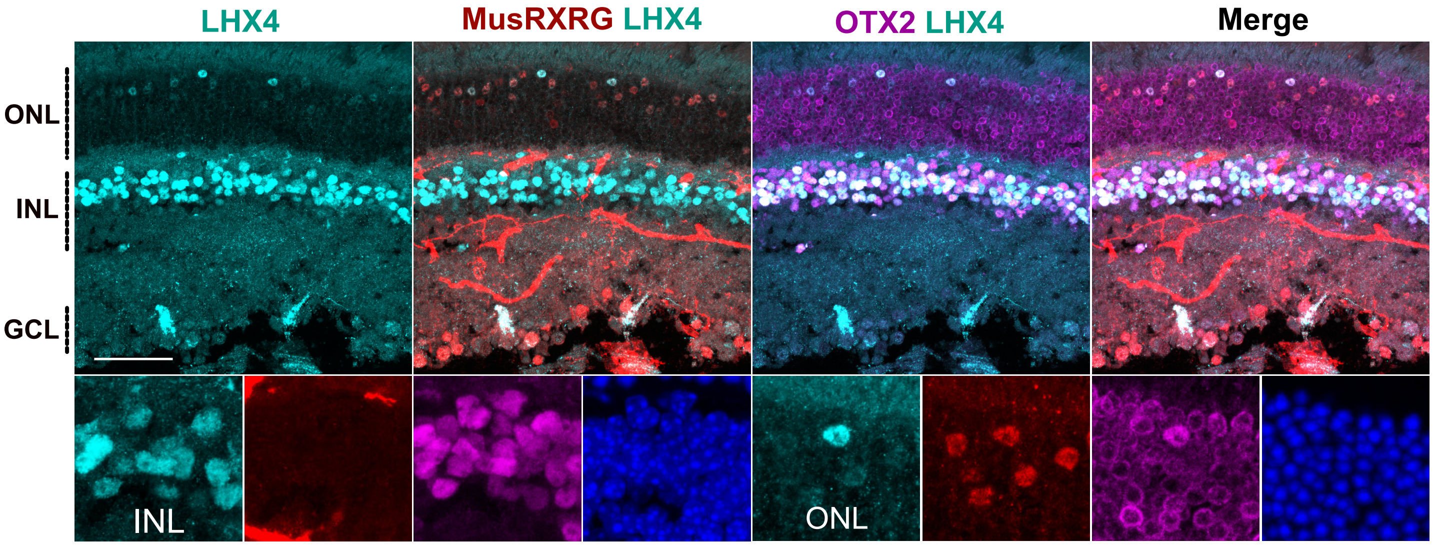


Supplemental Figure 3 - **LHX4 is expressed in cone bipolar cells and a subpopulation of cones in the adult mouse retina.**

Cross-section of a P27 mouse retina imaged for LHX4, RXRG, and OTX2. Higher magnification panels in the ONL and INL as specified. Large panels are maximum intensity projection Z-stack and small panels are single planes of the same Z-stack. Scale bar represents 50 μm

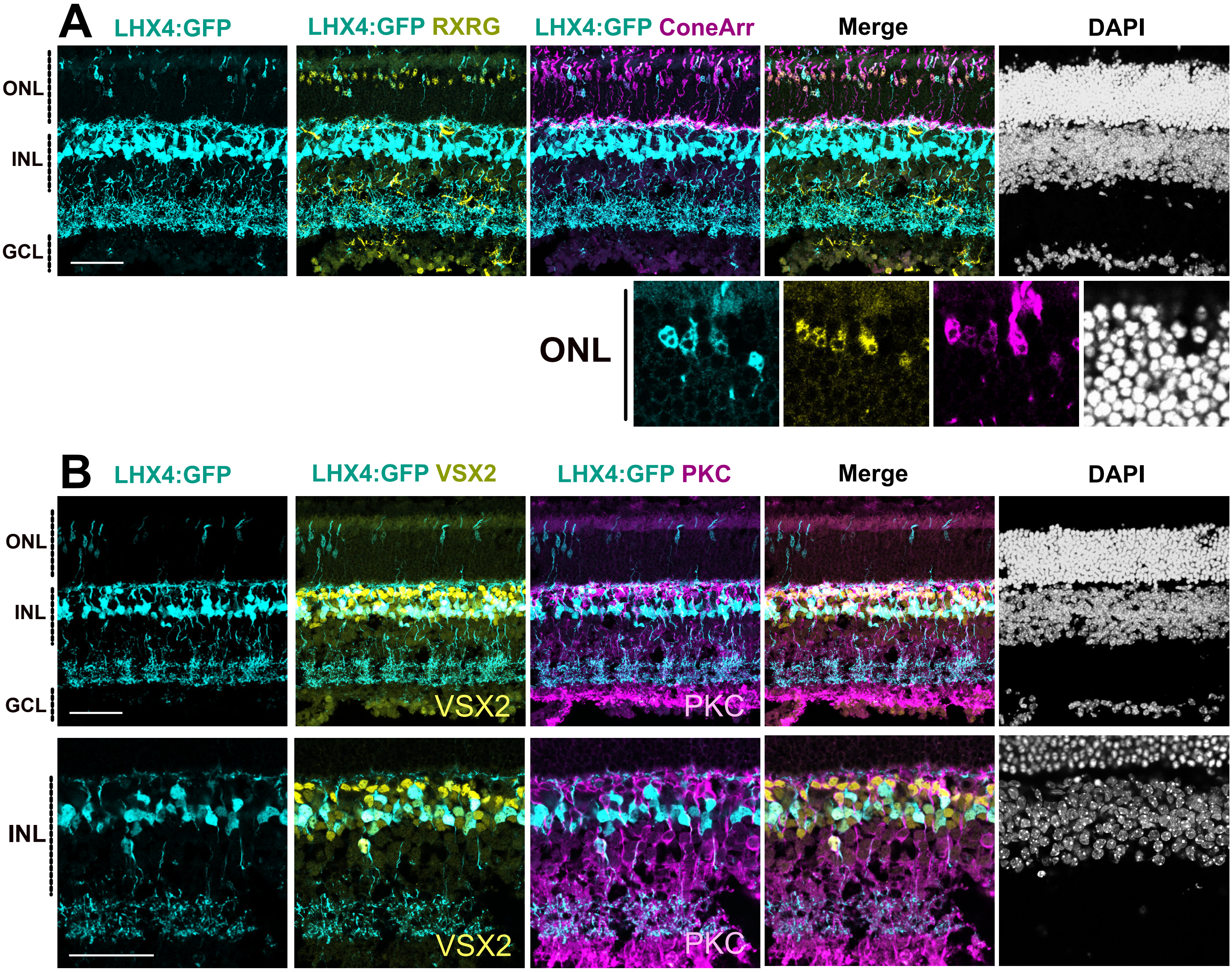


Supplemental Figure 4 - **LHX4-GFP reporter is active in cone bipolar cells and a subset of cones in the adult mouse retina.**

Cross-section of P27 mouse retinas imaged for EGFP, RXRG, and Cone Arrestin in (A), and EGFP, VSX2 and PKC in (B). Higher magnification panels in (A) show a single z-plane in the ONL. Large panels are maximum intensity projections Z-stacks and small panels are single planes of the same Z-stacks. Scale bar represents 50 µm

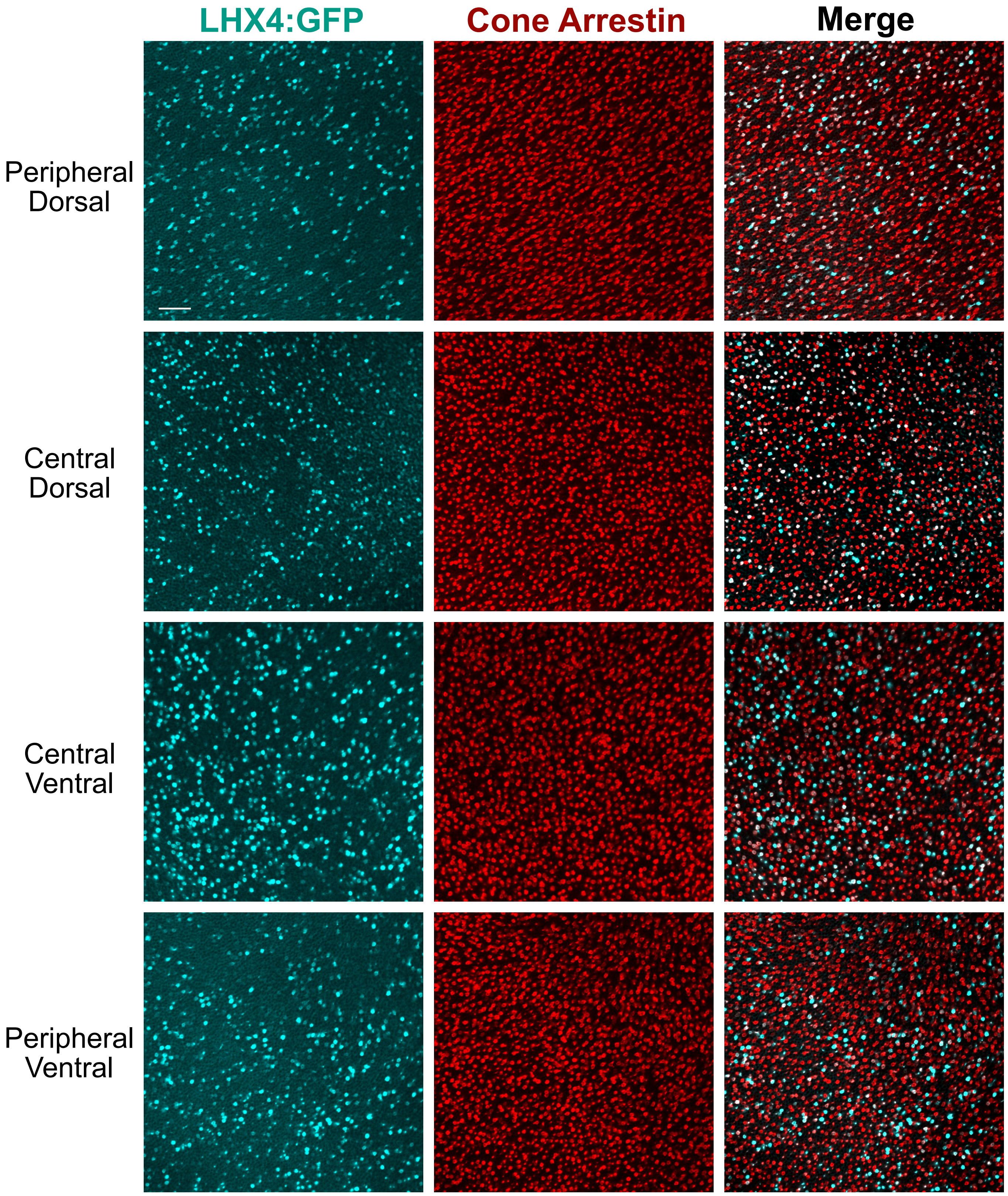


Supplemental Figure 5 - **LHX4-GFP reporter in the adult is active is not subject to dorsal-ventral gradient.**

Whole mount of a P27 LHX4-GFP mouse retina imaged for EGFP and Cone Arrestin imaged at the ONL. Scale bar represents 50 μm

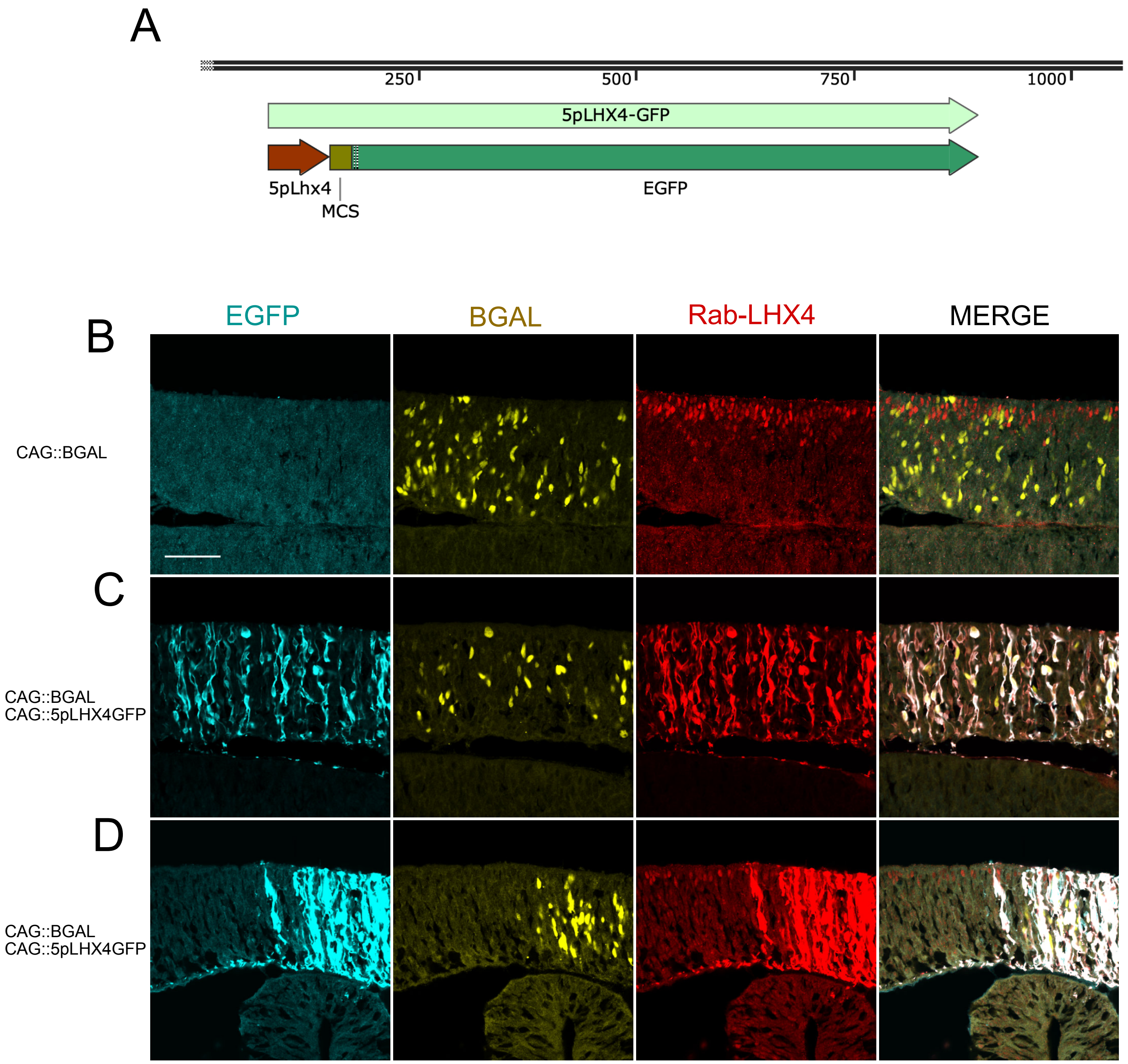


Supplemental Figure 6 - **LHX4-GFP insertion retains part of the LHX4 5' coding sequence and produces a N-terminusLHX4-GFP fusion.**

(A) Schematic of coding region amplified from LHX4-GFP locus containing a portion of the N-terminus of LHX4.

(B-D) Cross-section of chicken retinas after electroporation and ~20hr incubation. Retinas are imaged for EGFP, RbLHX4 and Bgal. In the case of (C), imaged at the edge of the electroporated area with increased exposure to show endogenous LHX4 signal. Scale bar represents 50 μm

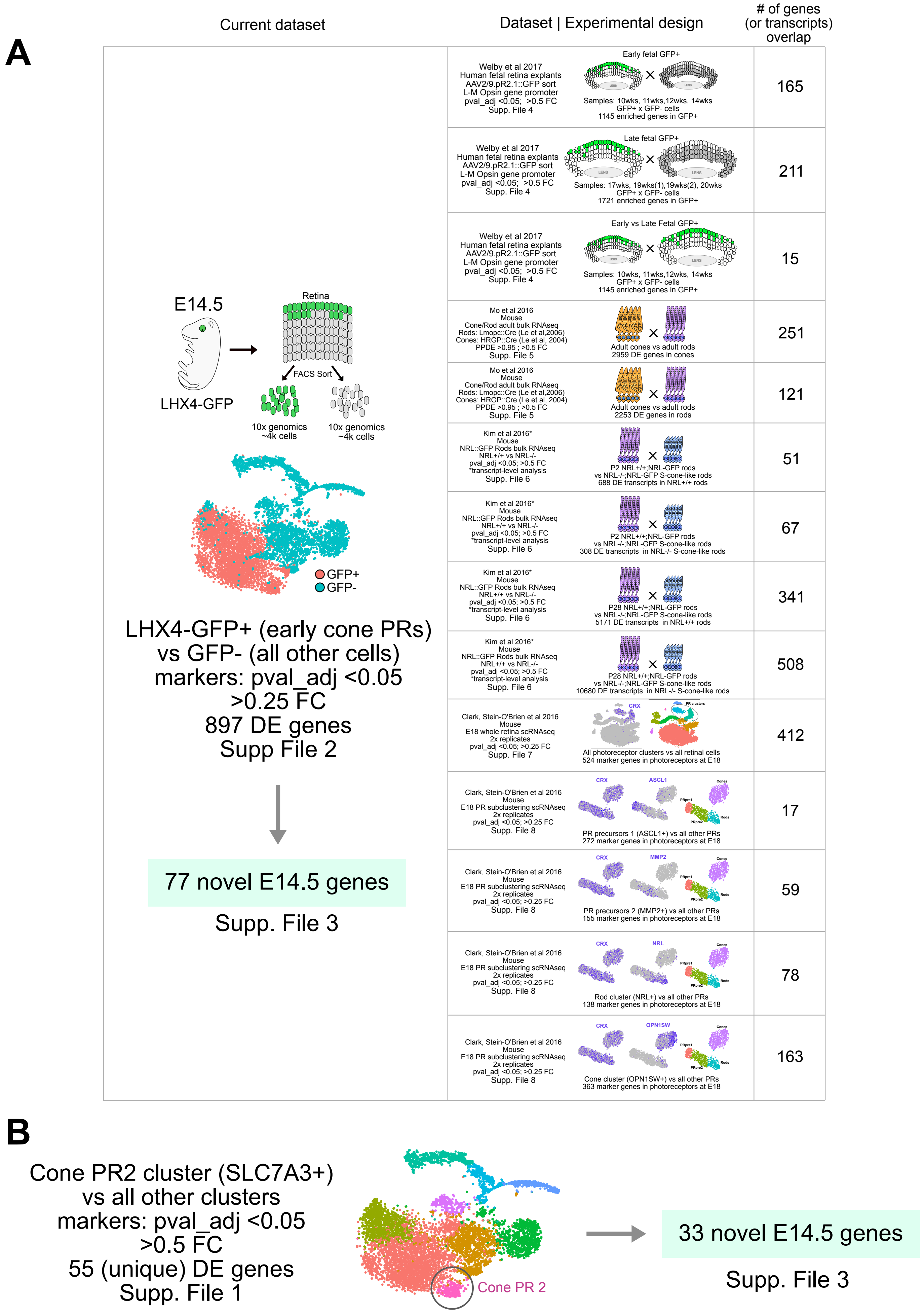


Supplemental Figure 7 - **Summary of datasets compared with E14.5 cone-enriched genes from LHX4-GFP current datasets.**

A) Table describing the datasets used for comparison and the amount of overlapping genes with E14.5 cone-enriched genes. For each report (Welby, Mo, Kim and Clark), datasets are divided in subsections depending on enrichment in a particular cell type, as denoted.

B) Summary of comparison with Cone 2 cluster enriched genes.

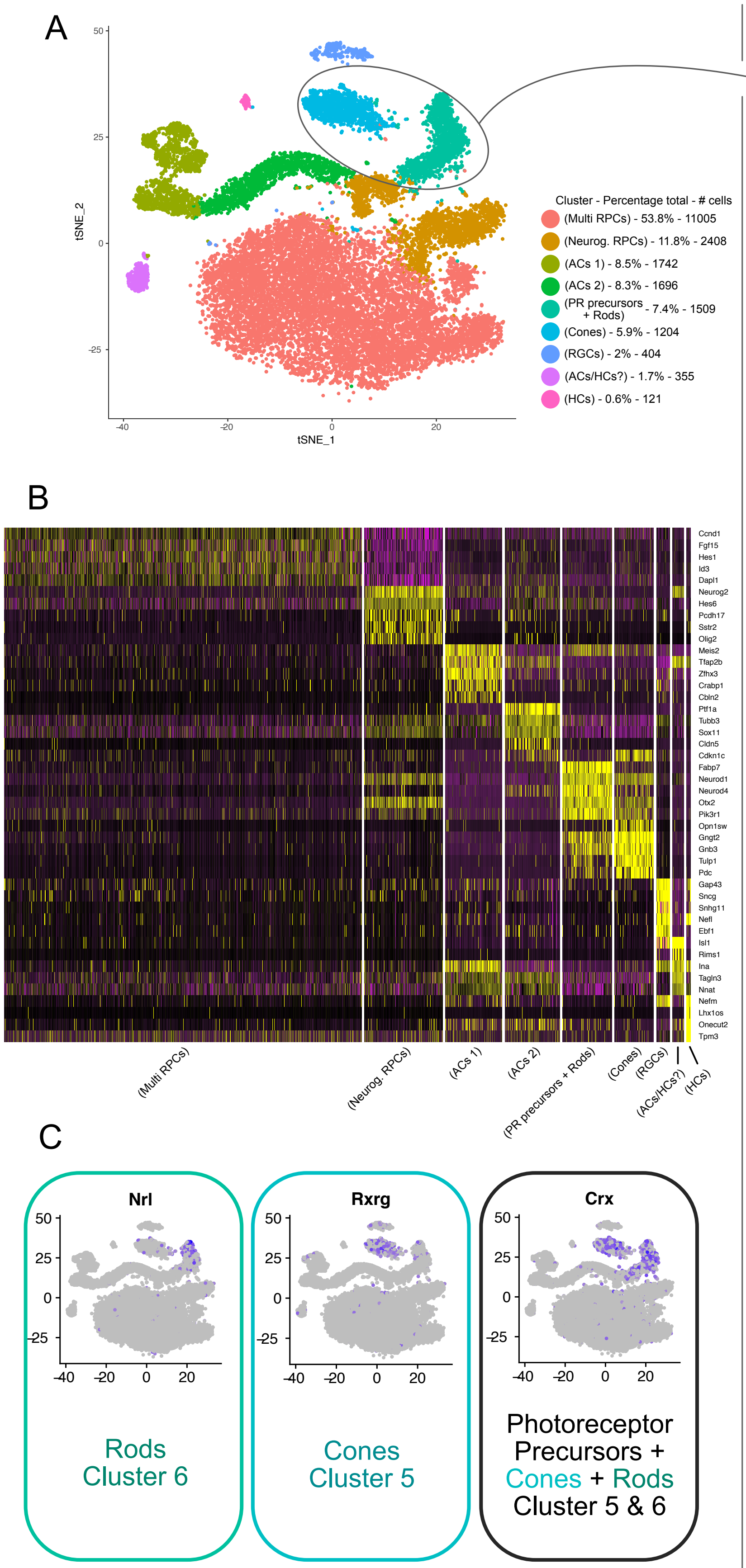


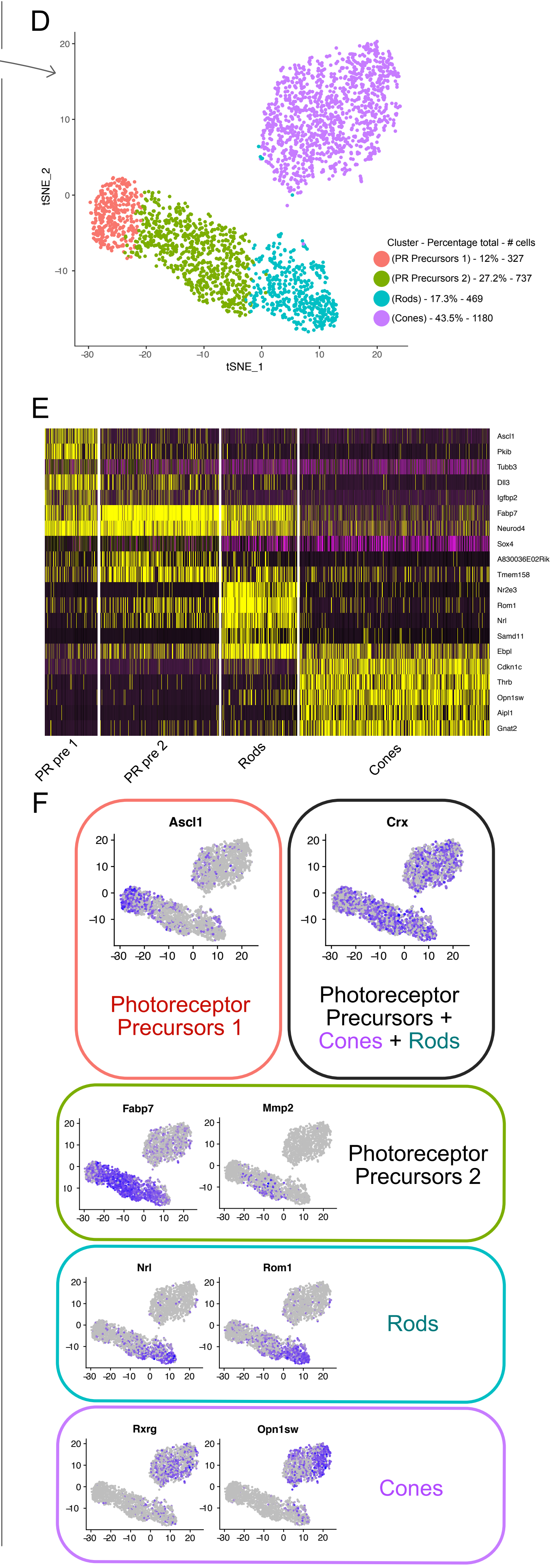


Supp Fig 8 - Clark Blackshaw e18 - **Unsupervised analysis of e18 time point from Clark & O'Brien et al 2018.**

A. TSNE plot of two replicates of e18 mouse whole retinas. Clusters are assigned according to markers shown in B and C. Highlighted are the photoreceptor clusters selected for further subclustering and analysis in D, E and F.

B. Heatmap of top 5 markers for each cluster.

C. Photoreceptor transcript expression projected onto the TSNE plot, as used for selecting clusters for further analysis.

D. TSNE plot of e18 photoreceptors after sub-clustering analysis. Clusters are assigned according to markers shown in E and F.

E. Heatmap of top 5 markers for each photoreceptor cluster.

F. Photoreceptor transcript expression projected onto the TSNE plot, as used for cluster assignment..
